# Accelerated and Reproducible Fiji for image processing using GPUs on the cloud

**DOI:** 10.1101/2022.07.15.500283

**Authors:** Ling-Hong Hung, Evan Straw, Zachary Colburn, Ka Yee Yeung

## Abstract

**Summary:** Graphical processing units can greatly accelerate image processing but adoption has been hampered by the need for specialized hardware and software. The cloud offers inexpensive on-demand instances that can be pre-configured with the necessary software. Specifically, we use the Biodepot-workflow-builder (Bwb) to deploy a containerized version of Fiji that includes the CLIJ package to use GPUs on the cloud. In addition, we provide an Amazon Machine Image (AMI) with the correct drivers and Docker images pre-loaded. We demonstrate the portability and reproducibility of the platform by deploying an interactive Fiji/CLIJ workflow on both Amazon Web Services and IBM cloud. The workflows produce identical results while providing a 29-fold reduction in execution time.

## 1 Introduction

Graphical processing units (GPUs) are specialized hardware that parallelize and accelerate common image manipulations, such as those used to render and display real-time graphics in computer games. They can also be used for to accelerate non-graphical computationally intensive tasks. Deep learning applications in particular, have benefited from the use of GPUs. However, With the exception of deep-learning, GPUs have not been widely used in open-source image processing applications. Recently, Haase *et al*. developed the CLIJ platform that adds GPU support for the Fiji/ImageJ software suite (Schindelin *et al*., 2012). Fiji is a popular open-source software suite that extends the basic image processing capabilities of ImageJ (Schneider *et al*., 2012) with an expansive set of plugins. Fiji also supports ImageJ macros, which can record and automate a series of processing commands. CLIJ leverages the Open Computer Language (OpenCL) framework to port image processing tasks to the GPU with significant speedups (Haase *et al*., 2020).

In this manuscript, we use the Biodepot-workflow-builder (Bwb) platform (Hung *et al*., 2019) to deploy a containerized Fiji/CLIJ and Jupyter workflow onto both Amazon Web Services and IBM Cloud.Container images are automatically downloaded to provide the necessary libraries and executables. Lower level GPU drivers that are not managed by the containers are provided in a pre-configured virtual machine image to further facilitate deployment. This is a portable and accessible methodology that integrates GPU-enabled image processing with other analytical tools to creates fast, interactive and reproducible multi-step analytical workflows.

## 2 Motivation

Using a GPU for accelerated processing can pose several challenges. GPUs are costly, may be difficult to obtain, have high power consumption and require tspecialized software, drivers and libraries. For many users, the cost and effort of configuring GPU software and maintaining a dedicated GPU server may be difficult to justify in spite of the reduced execution time. Our goal is to provide an open-source methodology that allows casual users to utilize GPUs to process their image data. We leverage the cloud to provide on-demand access to a high-performance GPU that incurs charges only when the server is used. Dependencies, libraries and drivers can be provided using downloadable software containers and preconfigured virtual servers. This greatly simplifies the onerous process of reproducibly deploying GPU workflows on the cloud.

## 3 Implementation

The Biodepot-workflow-builder (Bwb) (Hung *et al*., 2019) is a containerized platform for the creation, deployment and execution of analytical workflows consisting of graphical modules or widgets. Each widget manages its own Docker software container to encapsulate the executable or script and its software dependencies. Bwb modules use X11 commands to draw to a framebuffer (Hung *et al*., 2016) which is then viewed using an external VNC client or a VNC client (noVNC) implemented in the browser (Mittal *et al*., 2017). The result is a portable interactive desktop environment for containerized graphical applications that can be deployed on cloud servers or locally on laptops and desktops.

In this paper, we have used Bwb to add a graphical interface to CLIJ which allows users to configure CLIJ parameters using a form-based user interface. The Bwb Fiji/CLIJ widget uses the user inputs to generate a shell command that executes Fiji. Execution takes place inside a Docker container image containing Fiji with the CLIJ plugin, OpenCL and NVIDIA libraries. Bwb usually only require the installation of Docker to start Bwb, a VNC viewer or browser to connect to Bwb and interact with the graphical desktop. The executables and software dependencies are automatically and reproducibly installed by Bwb which pulls the required Docker containers from our public repository. However, communicating with GPUs also requires low-level drivers that are not managed by containers but by the underlying operating system kernel. The user must install the correct version of drivers that match the version of the GPU software in the containers. Our containers are tightly version controlled and we include the GPU software version in the naming convention to allow the user to determine driver versions required. For cloud deployments, we provide virtual machine images in which Docker, the container images and drivers are all pre-loaded. Starting a cloud instance using this machine image eliminates the need for manual installation of software. Finally, users can also use their own existing Fiji installations by specifying the installation directory.

## 4 Results

To demonstrate the utility and performance of our implementation of CLIJ, we created a Bwb workflow (Figure 1) which downloads the image dataset, runs a Fiji macro to process the data on GPU and CPU, uses Jupyter to visualize execution times and compares any differences in the results. The workflow can be run automatically or interactively, with the user being able to stop and adjust parameters and restart at any step. Light-sheet microscopy test data, segmentation macros and Jupyter notebooks that make up the workflow were obtained from the original CLIJ paper (Haase *et al*., 2020). The macros/notebooks were modified from the original to produce histograms of the average execution time and to use the file paths specified by Bwb workflow. A short demonstration video of the running workflow is available at https://youtu.be/3nPYKpV04bc.

**Figure 1:**
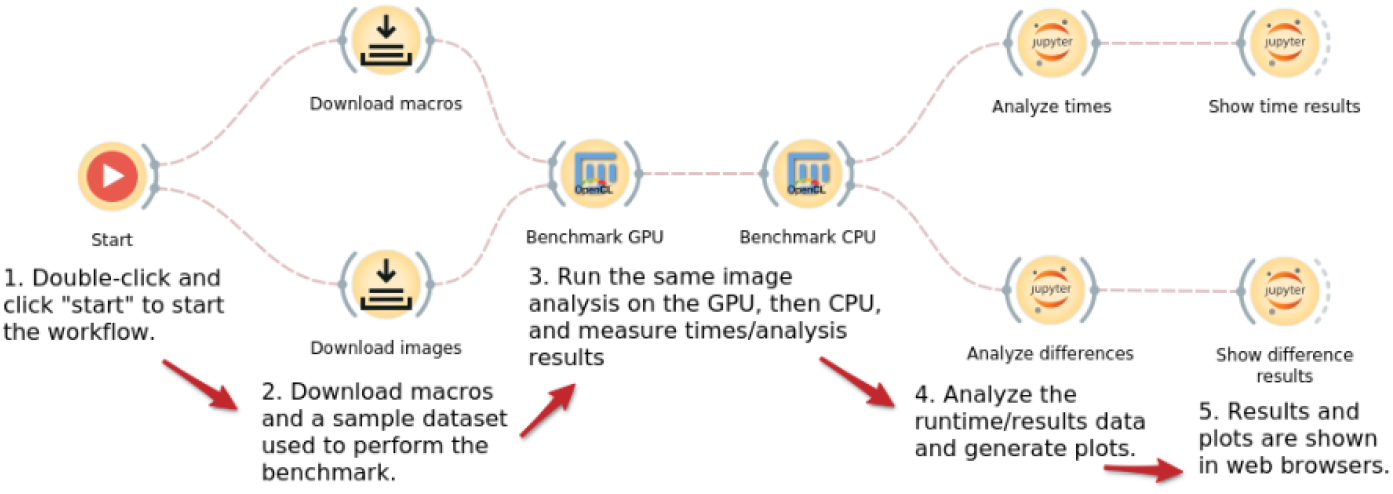
CLIJ example workflow implemented in the Biodepot-workflowbuilder. This workflow includes both the GPU and CPU versions of Fiji to benchmark and compare performance. Since both the GPU and CPU enabled Fiji widgets are encapsulated in their own Docker containers, the user can easily build modular containerized workflows that require different computing environments.

The workflow was benchmarked on an AWS EC2 g4dn.2xlarge instance (8 vCPUs, 32GB of RAM, and an NVIDIA Tesla GPU with 16GB of VRAM) and on an IBM Cloud `gx2-8×64×1v100` instance (with 8 vCPUs, 64GB of RAM, and an NVIDIA Tesla V100 GPU with 16GB of VRAM). We found that the entire processing of one image (including I/O operations) on the GPU in the CLIJ benchmark runs on average about 13 times faster than the processing done using the CPU(see Figure 2). Excluding I/O operations, the GPU workflow is 29 times faster than the CPU workflow. Additionally, although there were small differences between the results of the GPU and CPU analyses, the results between the two GPU analyses performed on AWS and IBM Cloud were identical (see Figure 3).

**Figure 2:**
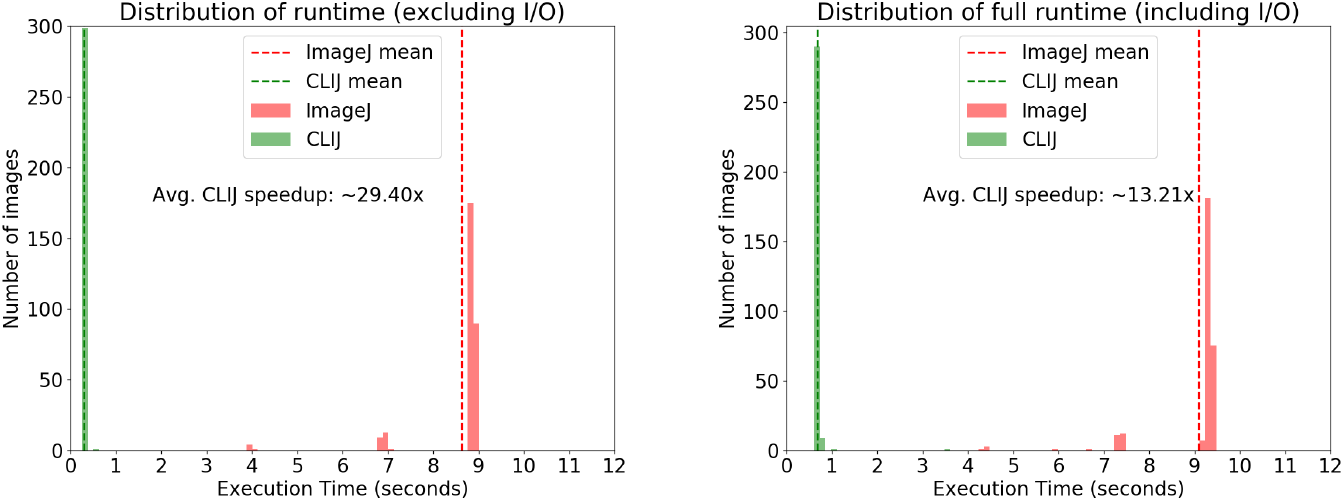
Distribution of the time taken to process one image in the GPU (on CLIJ) and CPU (on ImageJ) benchmarks, across 300 images. The left histogram shows the execution time with I/O operations excluded while the right histogram shows the execution time including I/O operations.

**Figure 3:**
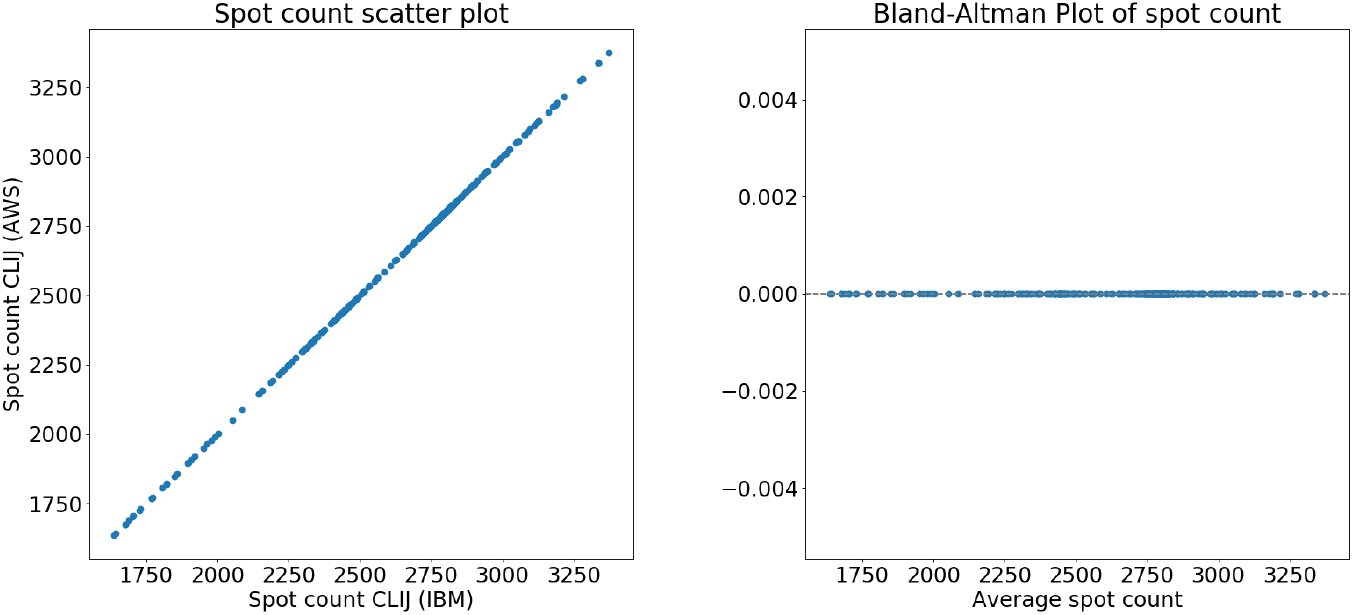
Results from running the workflow on IBM Cloud versus running it on AWS. The left panel shows a scatter plot comparing the execution time of each image using CLIJ deployed on AWS versus IBM Cloud. The right panel shows a Bland-Altman plot (difference plot) between the AWS execution time and IBM execution time.

## 5 Discussion and conclusions

Our performance benchmark demonstrates substantial speedup (29.40x excluding I/O operations, 13.21x including I/O operations) using our implementation of Fiji/CLIJ. The AWS and IBM cloud servers had identical GPUs and as expected, the execution times were very similar. AWS and IBM use different hypervisors to virtualize the kernel which could potentially affect the driver level interactions with the GPU. However, both GPU platforms produced identical results, which highlights the portability and reproducibility of the containerized Bwb workflows. There were some small differences between GPUs and CPU results. This was also observed in the original CLIJ paper and is not surprising given the differences between GPU and CPU algorithm implementations and data precision.

There are some limitations of the Bwb Fiji/CLIJ methodology. One is that the user must manage data transfer to and from the cloud. There are many tools for this including sftp, sshfs, as well as vendor specific tools. We are also actively working on a built-in transfer tool for Bwb. A second limitation involves hardware incompatibilities that are not masked by Docker containers. Although CLIJ uses open-source OpenCL libraries supported by both NVIDIA and AMD GPUs, the Docker container does not recognize AMD devices as GPUs. Also, there ARM devices such as M1/M2 Macs are problematic for Fiji as most tools, libraries and Docker containers are based on x86 architectures. We are working to overcome these limitations in future releases of Bwb. However, cloud usage is not unduly affected as the major cloud vendors all have instances with x86 CPUs and NVIDIA GPUs.

To summarize, we present an open-source, accelerated and containerized image processing platform that is easily deployed on GPU servers on different cloud platforms. We believe that our platform will make GPU-enabled image processing more accessible to users in an on-demand cloud computing model.

## Acknowledgements

LHH and KYY are supported by NIH grant R01GM126019. LHH, ES, ZC and KYY are supported by NCI SBIR contract 75N91021C00022. We would also like to thank Amazon Web Services and IBM Cloud for cloud credits. The content is solely the responsibility of the authors, and does not necessarily represent the official views of the National Institutes of Health.

LHH and KYY also have equity interest in Biodepot LLC, which receives compensation from NCI SBIR contract numbers 75N91020C00009 and 75N91021C00022.

## Footnotes

0 Adapted from the versions given in the Supplementary Materials section of the CLIJ paper (Haase *et al*., 2020)

## Notes

### Competing Interest Statement

LHH and KYY have equity interest in Biodepot LLC, which receives compensation from NCI SBIR contract numbers 75N91020C00009 and 75N91021C00022.

